# Epigenetic Liquid Biopsy Enables Universal Mutation-Agnostic Molecular Surveillance for High-Risk Neuroblastoma

**DOI:** 10.64898/2026.01.15.699217

**Authors:** Nurit Gal-Mark, Assaf Grunwald, Valid Gahramanov, Michal Hameiri-Grossman, Elena Shinderman-Maman, Dafna Gaas, Keren Shichrur, Eva Chausky Barzakh, Aviv Sever, Shirah Amar, Shifra Ash, Yehudit Birger, Shai Izraeli, Yuval Ebenstein, Esther R. Berko

## Abstract

Pediatric solid tumors present a challenge for precision oncology as most tumors harbor low mutational burden, limiting the applicability of standard mutation-based liquid biopsy approaches for molecular monitoring. To overcome this, we developed a mutation-independent liquid biopsy framework based on Oxford Nanopore Technology sequencing, utilizing robust DNA methylation-based biomarkers for detection and surveillance of high-risk neuroblastoma (HR-NBL), a common pediatric solid tumor. Comparative analysis of tumor-derived DNA methylation profiles against a comprehensive atlas of normal human cell types identified 72 NBL-specific differentially methylated regions (termed meNBLs). Integration of 25 selected meNBLs into the methylation atlas enabled quantitative deconvolution of NBL-derived cfDNA, outperforming copy number and mutation-based estimates. NBL-specific blood plasma-derived cfDNA was detected at diagnosis but not in healthy controls, absent during remission, and markedly elevated at relapse, even before clinical detection of disease recurrence. These findings establish methylation-based deconvolution as a robust, mutation-independent approach for cfDNA monitoring in HR-NBL.

## Introduction

Neuroblastoma (NBL), a malignancy of the developing sympathetic nervous system, is the most common extracranial solid tumor of early childhood. The disease is highly heterogeneous, with clinical behavior ranging from spontaneous regression to aggressive therapy-resistant progression (1). Despite extensive and highly morbid treatments, event-free survival in high risk (HR) NBL remains ∼50%, and outcomes following relapse are dismal.

Like other pediatric solid tumors, HR-NBL is characterized by a relative paucity of single nucleotide variants (SNVs) at diagnosis (2). While several genomic alterations associated with aggressive disease have been described, most are not amenable to standardized SNV-based molecular tracking. *MYCN* amplification is a well-established driver of HR-NBL and confers aggressive biology. Activating aberrations in the Anaplastic Lymphoma Kinase (*ALK*) gene occur in ∼20% of patients, portend inferior prognosis, and remain the only currently tractable target in this disease (3, 4). Additional alterations involving the TERT promoter locus, ATRX, and the RAS-MAPK pathway occur in subsets of tumors and are associated with inferior outcomes (5). However, the majority of HR-NBL tumors lack recurrent, clonal SNVs suitable for longitudinal molecular monitoring and are instead characterized only by classic segmental chromosomal aberrations (SCA) (6).

Liquid biopsy analysis of circulating tumor DNA (ctDNA) has emerged as a powerful, minimally invasive approach to capture tumor genetic heterogeneity, monitor disease burden, and detect clonal evolution in HR-NBL (7, 8). While ultra-deep targeted sequencing panels and mutation-specific PCR assays have demonstrated clinical utility (7–11), their application is limited to approximately 50% of HR-NBL patients whose tumors harbor trackable driver oncogenic alterations. This underscores the unmet need for universal mutation-independent biomarkers for liquid biopsy-based monitoring across the full HR-NBL population.

DNA methylation patterns are highly cell-type-specific, stable across cell divisions, and universally present, making them compelling candidates for mutation-agnostic liquid biopsy approaches. Prior methylation studies in NBL have largely focused on identifying prognostic signatures by comparing high-risk tumors to low-risk disease or normal tissues (12–14). While informative, these approaches were not designed to enable sensitive and specific detection of tumor-derived DNA within the complex mixture of cell-free DNA (cfDNA) and are utilized to interrogate only a subset of genomic loci. Here, we adopt an alternative strategy. Building on methylation-based deconvolution frameworks established by Dor and co. (15, 16), we identify differentially methylated regions (DMRs) that uniquely distinguish NBL from all other normal human cell types represented in the comprehensive methylation atlas. Using Oxford Nanopore Technologies (ONT) sequencing, which enables direct methylome profiling from native DNA while simultaneously capturing SNV and copy number alterations (CNAs), we integrate HR-NBL as a distinct entity within the reference atlas. We demonstrate that this approach enables the sensitive and quantitative detection of NBL-derived cfDNA across different disease states, supporting methylation-based deconvolution as a robust and universal framework for longitudinal monitoring of NBL.

## Results

### Identification of novel NBL-specific epigenetic markers

To identify robust NBL DMRs we performed ONT sequencing on seven HR-NBL tumor samples, achieving a median coverage of 32X genome-wide. The seven tumor samples were selected to represent the major molecular subtypes and genetic alterations characteristic of HR-NBL (Supplementary Table S1). ONT sequencing profiled concurrent genome-wide genetic (SNV and CNV) and epigenetic (CpG methylation) data from each sample (Figure 1A). We accurately detected the known clinically relevant SNVs and SCAs in each patient tumor, as previously identified in clinical testing with multiple disparate diagnostic approaches (Figure 1B). Additionally, the long-read ONT approach enabled identification of novel structural variants (SVs) not seen in clinical testing panels (Supplementary Table S1). Genome-wide DNA methylation profiling revealed globally reduced levels of DNA methylation in NBL tumor genomic DNA compared with control mixed PBL DNA samples (Supplementary Figure S1A), consistent with the well-established global hypomethylation observed across many cancer types (17). Hierarchical clustering and principal component analysis (PCA) based on global methylation patterns clearly distinguished NBL tumors from PBL samples and further subdivided NBL tumors according to molecular subtypes, in agreement with previously reported classifications (18, 19) (Figure 1C and Supplementary Figure S1B).

**Figure 1.**
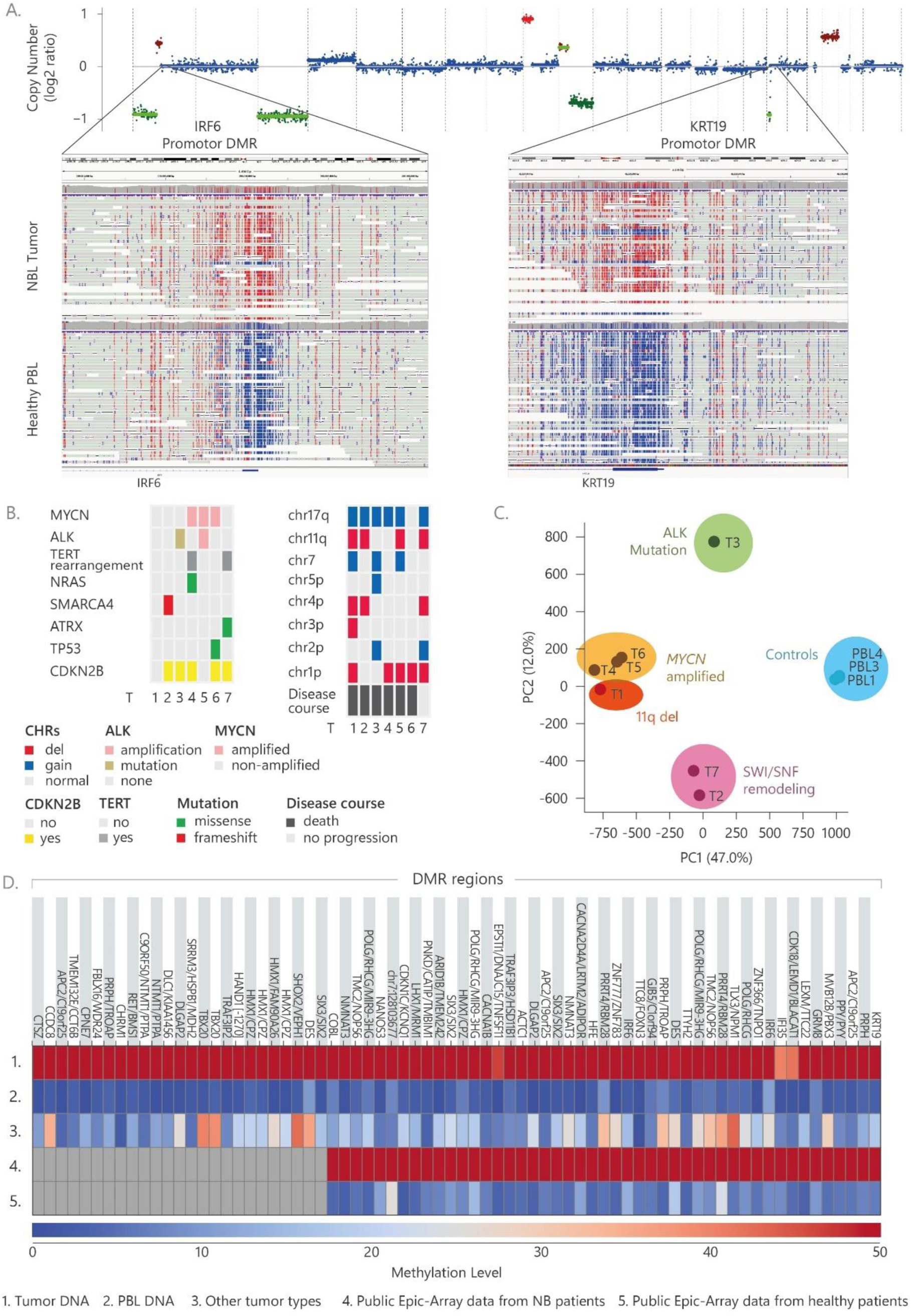
Integrated genetic and epigenetic profiling of NBL using whole-genome ONT sequencing. **(A)** Whole-genome ONT sequencing of tumor DNA provides simultaneous genetic and epigenetic information, illustrated here for patient T6. The data enable detection of large-scale genetic alterations, shown as a CNV profile in the upper panel, and identification of DMRs relative to control samples in the lower panel. The lower panel highlights two representative promoter-associated DMRs: the newly identified IRF6 promoter on chromosome 1 and the previously reported KRT19 promoter on chromosome 17. Individual native DNA molecules aligned to each locus are displayed, with CpG sites colored red (methylated) or blue (unmethylated). **B)** Oncoplot: Major genetic alterations associated with NBL were identified by whole-genome ONT sequencing and summarized across the seven sequenced tumors. The oncoplot displays clinically relevant genomic events, including gene amplifications, mutations, deletions, chromosomal copy number aberrations, and clinical disease course annotations. **(C)** PCA of genome-wide DNA methylation profiles from the seven NBL tumors reveals clustering according to key genomic alterations, including MYCN amplification and 11q deletion, ALK mutation, and SWI/SNF remodeling, with control samples forming a distinct cluster. **(D)** NBL DMR heatmap: DNA methylation data from the seven NBL tumors were compared against a reference atlas comprising methylomes from 39 healthy tissue types to identify pan-NBL–specific DMRs. Methylation levels across the 72 identified DMRs are shown in the heatmap, with colors indicating average methylation levels (red, high methylation; blue, low methylation); values represent group-level averages for each column, and arranged in rows representing from top to down: the seven sequenced NBL tumors, peripheral blood leukocyte (PBL) samples as healthy controls, public methylation data from other advanced tumor types, and previously published NBL and healthy control methylomes generated using Illumina EPIC arrays. NBL DMRs not interrogated by the EPIC array are shown in gray.

Leveraging the methylation atlas of normal human cell types (15), we identified 72 novel NBL-specific DMRs, herein referred to as meNBLs (Figure 1D, see methods for DMR selection criteria). Notably, although we observed global hypomethylation in NBL tumor genomes, the NBL-specific DMRs were predominantly hypermethylated relative to other tissues in the human methylation atlas (Supplementary Table S3). The existence of highly methylated regions within a globally hypomethylated background has likewise been reported in previous studies of NBL (20).

Gene ontology analysis revealed that genes associated with these NBL-specific DMRs were significantly enriched in biological processes related to embryonic organ development, cognition, and neuronal maturation, consistent with the expected lineage and developmental origin of NBL (Supplementary Figure S3 and Supplementary Table S4). The 72 meNBLs identified were found to be distributed across various genomic regions, including promoters, gene bodies, 3′ untranslated regions (3′ UTRs), and intergenic regions, with several mapping to long non-coding RNAs (Supplementary Figure S4). In addition to established DNA methylation biomarkers in NBL, such as PRPH and KRT19 (12, 13), several novel candidates were identified. These include transcription factors such as HAND1, SIX3, SHOX2, HMX1, and IRF6 and TBX20, many of which are known to play roles in embryonic and neuronal development. Additionally, genes encoding calcium and potassium channels, CACNA1B and CACNA2D4, were also identified, all of which are involved in neuronal signaling. The involvement of these developmental regulators aligns with the embryonic origin of NBL and highlights pathways whose dysregulation may contribute to aberrant differentiation and tumorigenesis.

To further evaluate the robustness of our meNBL panel, we used publicly available EPIC array data, comprising 205 NBL tumors and 152 healthy controls (21). Many of the EPIC array features overlap with our meNBL panel and display similar distinct methylation level differences between NBL and controls (Figure 1D). Notably, ∼30% of our meNBLs (highlighted in grey in Figure 1D) are not covered by the EPIC array, highlighting the potential of our approach in discovering novel diagnostic biomarkers and potential therapeutic targets that are inaccessible to EPIC arrays. To assess the specificity of meNBLs, we compared them to the methylation profiles of tumors from an independent ONT-sequencing cancer dataset (22) (Supplementary Table S9). This benchmarking analysis confirmed that our selected meNBLs contain both DMRs that are a) unique to NBL, reflecting the neuronal cell of origin of NBL when compared to other malignancies or, b) shared DMRs with other cancer types. The combination of these markers enables accurate detection of this peripheral nerve malignancy by confirming both its carcinogenic nature and its neuronal origin (Figure 1D).

### Application of meNBLs to liquid biopsy-based disease detection

To assess whether meNBLs can be detected in cfDNA and utilized for liquid biopsy, we performed ONT sequencing on cfDNA obtained at diagnosis from 12 HR-NBL patients. Five cfDNA samples were matched to sequenced tumor tissues, and an additional seven cfDNA samples served as an independent validation cohort. Additionally, we sequenced cfDNA from 8 healthy individuals to serve as controls (median coverage of 2.8x, Supplementary Table S2). Importantly, analysis of meNBL DMRs showed concordance across all loci in all HR-NBL patients with no significant differences in methylation levels between the matched and independent cohorts were observed (t-test) (Figure 2A). These findings confirm that the selected markers are not restricted to the initial patients whose tumor was used for marker discovery supporting their broader applicability as NBL-specific biomarkers.

**Figure 2.**
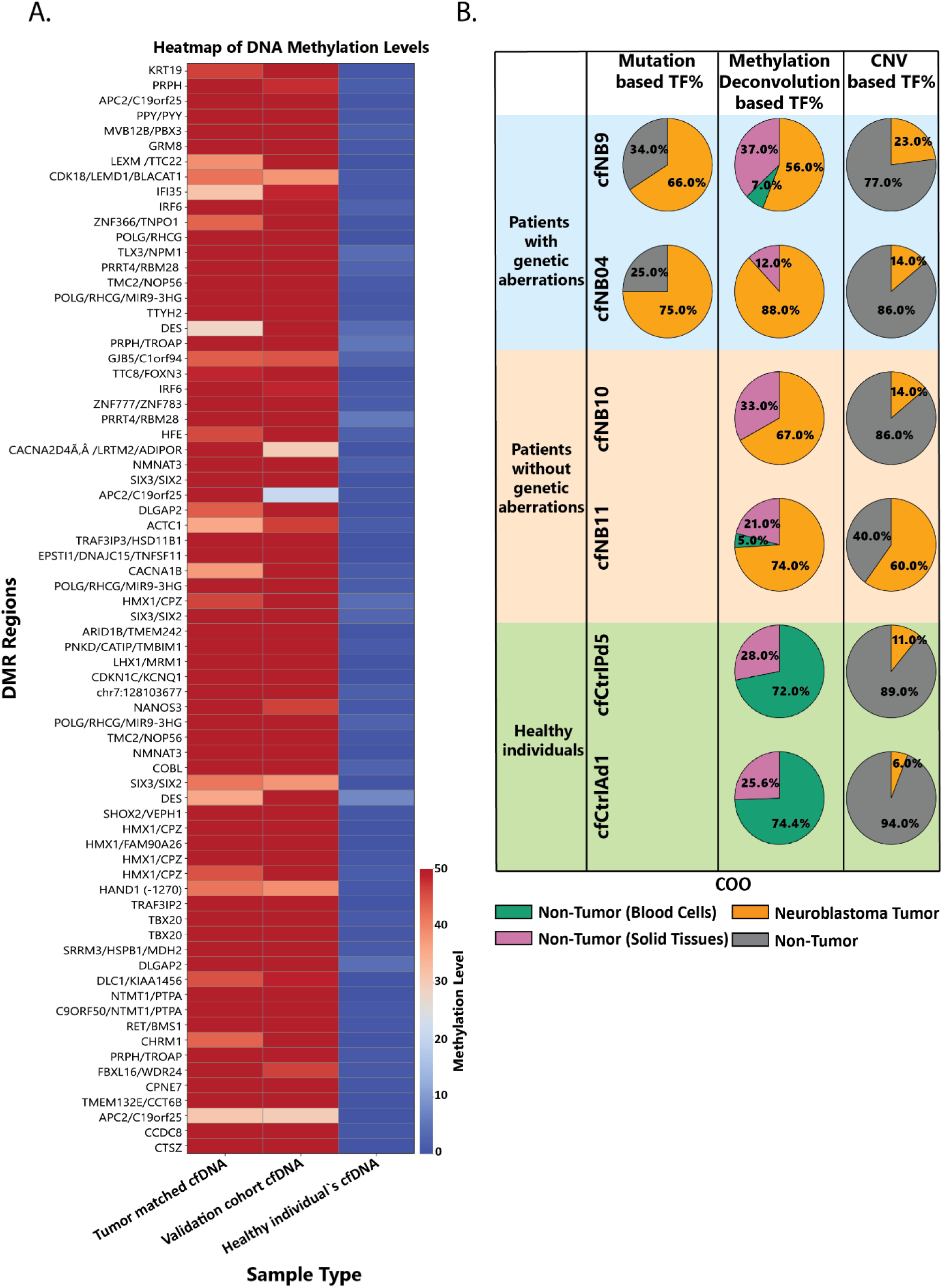
Detection of NBL markers in cfDNA samples. **(A)** Heatmap showing average DNA methylation levels across the 72 identified DMRs. Rows correspond to individual DMRs and columns represent cfDNA sample groups: matched cfDNA from patients with sequenced tumor tissue, cfDNA from an independent validation cohort of NBL patients, and cfDNA from healthy individuals. Colors indicate mean methylation levels (red, high methylation; blue, low methylation). Values represent group-level averages across samples within each category. **(B)** Tumor fraction (TF%) estimates in cfDNA for representative individuals, stratified by clinical and genomic status, using three independent approaches. Columns show TF% estimation based on: (left) somatic mutation allele frequencies, applicable only to patients with identifiable tumor mutations; (middle) methylation-based deconvolution using our novel NBL markers; and (right) copy-number variation (CNV) based estimation using ichorCNA. Pie charts depict the relative contribution of tumor-derived cfDNA (orange-red) and non-tumor cfDNA fractions. Only the methylation-based approach further resolves the non-tumor fraction into cell of origin components.

To develop a quantitative NBL cfDNA assay, we leveraged a cell-of-origin deconvolution approach (15, 16). This method enables quantifying the contribution of different cell types to the total cfDNA content extracted from a given sample. We hypothesized that by defining NBL as an additional cell type in the human methylation atlas, we could detect NBL signatures that correlate with the existence and state of disease. As the atlas includes 25 markers per cell type, we selected 25 meNBLs from the 72 NBL-specific markers and incorporated them back into the atlas as a distinct NBL entity (see material and methods). We performed methylation-based deconvolution on all our cfDNA samples using our modified atlas. No NBL fraction was detected for the control cfDNA samples while varying amounts of NBL ctDNA were detected for all of the patient liquid biopsies (Figure 2). To validate the reliability of our tumor fraction quantification (TF%), we compared NBL-deconvolution estimates to those derived from the widely-used CNA-based analysis using IchorCNA, an algorithm designed for low-coverage sequencing samples (23). Additionally, for samples harboring somatic driver mutations, we also calculated SNV-based estimates (based on maximum variant allele frequency, with sufficient coverage). In approximately half of NBL cases, IchorCNA and methylation-based deconvolution yielded similar TF% estimates (Supplementary Table S7). Notable discrepancies highlighted the advantage of methylation-based deconvolution (Figure 2B): in healthy controls, IchorCNA detected significant background noise TF% (6–11%), whereas methylation-based deconvolution showed no sign of NBL (Figure 2B, green panel), demonstrating greater specificity. Two NBL cases where methylation TF% differed significantly from the IchorCNA assessment were found to have genetic drivers favoring the methylation based TF%. Sample cfNB4 (Figure 2B blue panel) displayed a NBL-deconvolution estimated 88% TF, in contrast to IchorCNA TF% of 14%. Based on cfDNA ONT sequencing data (8X coverage) of the known NRAS p.Gln61Lys mutation, a mutation-based TF% of 75% was calculated, closely aligning to methylation-based deconvolution scoring. Similarly, in cfDNA sample cfNB9, ichorCNA predicted 23% TF (despite prominent CNAs, see Supplementary Figure S5), while NBL-deconvolution estimated 56%. ddPCR confirmed an ALK p.Arg1275Gln mutation at variant allele fraction (VAF) of 32.9% yielding a mutation-based TF% of 65.8%, consistent with the methylation-based prediction. In addition, deconvolution enables estimation of contributions from other tissue types, providing further potential information. Taken together, these results demonstrate that methylation-based deconvolution is feasible in cfDNA and provides an accurate assessment of NBL contribution.

### meNBL markers for non-invasive monitoring of NBL disease progression

Based on our findings that Methylation-based liquid biopsy deconvolution using our meNBL DMRs detected NBL-derived cfDNA across all HR-NBL patient samples at varying proportions, while remaining undetectable (0%) in healthy donors (Figure 3A and Supplementary Table S7), we hypothesized that the approach may address the need for non-invasive molecular monitoring in NBL patients. To further validate NBL-deconvolution accuracy, we applied the algorithm to cfDNA samples collected from NBL patients during relapse and at molecularly confirmed remission time points (Supplementary Table S10). Methylation-based deconvolution analysis revealed the presence of NBL-specific cfDNA at diagnosis, its absence during confirmed remission time points, and marked elevation at relapse (Figure 3B).

**Figure 3.**
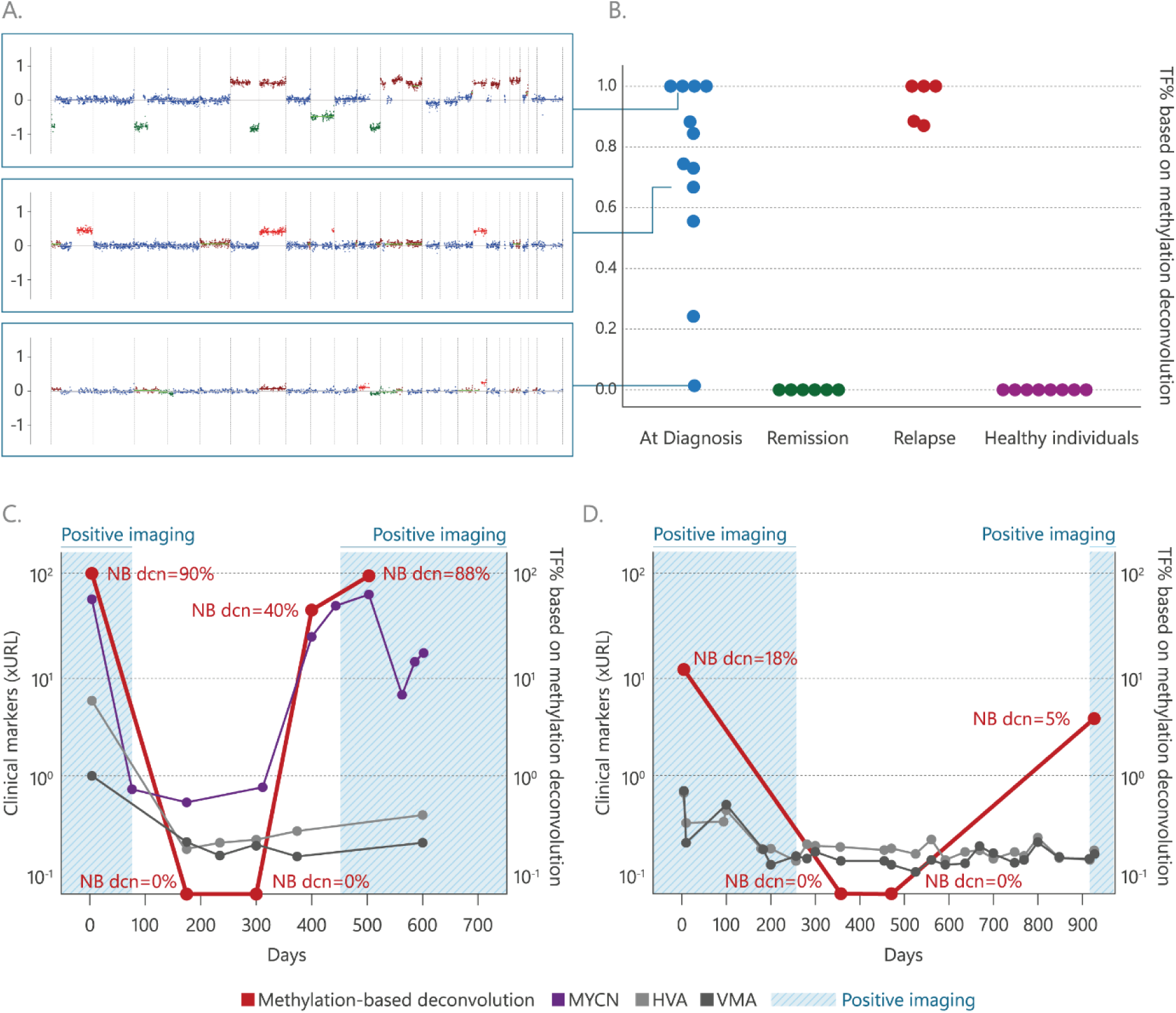
NBL methylation markers enable disease monitoring and longitudinal follow-up. **(A)** Representative copy-number variation (CNV) profiles derived from cfDNA samples collected at diagnosis from three NBL patients with varying tumor fractions (TF%). From top to bottom, examples are shown for patients with high, intermediate, and low TF%. **(B)** Tumor fraction (TF%) estimates obtained by methylation-based deconvolution using the 25 selected NBL markers, shown for cfDNA samples collected at different disease stages: diagnosis (N = 12), remission (N = 6), relapse (N = 5), and healthy individuals (N = 8). Each dot represents an individual cfDNA sample, and the Y-axis represents TF%. Arrows link representative TF% values in (B) to the corresponding CNV profiles shown in (A). **(C, D)** Longitudinal monitoring of NBL in two representative patients: a patient with MYCN amplification (C) and a patient without MYCN amplification (D). The x-axis denotes days from diagnosis. The left y-axis shows normalized values of established NBL disease markers, including MYCN copy number, and urinary catecholamine metabolites (HVA/VMA), as indicated in the legend. The right y-axis shows tumor fraction estimates derived from methylation-based deconvolution using the NBL marker panel. Blue shaded regions indicate time intervals during which gold standard meta-iodobenzylguanidine (MIBG) imaging confirmed active disease. In the MYCN-amplified case, methylation-based TF% tracking detected disease with sensitivity comparable to MYCN-based markers, while in the non-MYCN case, methylation-based detection mirrored imaging-based disease status.

To directly validate this approach against the clinical standard of care for both NBL subtypes, those with trackable genetic aberrations and those without, we present a time course of serial samples from one patient with known *MYCN* amplification (Figure 3C) and another with no detected NBL-associated genetic aberrations (Figure 3D). In both patients, meNBL liquid biopsy predictions were concordant with clinical status determined by imaging, revealing detectable NBL-derived cfDNA during active disease and none during known remission. In the *MYCN*-amplified patient, molecular assessment of disease presence was confirmed by ddPCR of cfDNA for *MYCN* amplification. In this patient, NBL-derived cfDNA was detectable with meNBL ∼100 days prior to clinically confirmed disease relapse, and consistent with plasma-based MYCN detection with ddPCR. These cases demonstrate that genomic molecular testing may outperform standard clinical assessment, including gold-standard urinary catecholamine biomarkers (VMA/HVA). Furthermore, due to its high accessibility via blood sampling, this liquid biopsy approach enables disease detection between imaging intervals, as demonstrated by longitudinal cfDNA profiling. Taken together, our results demonstrate that methylation-based deconvolution disease detection provides a robust and reliable estimator of NBL-derived cfDNA burden, and can provide specific and accurate information on disease for monitoring NBL progression and relapse.

## Discussion

Serial disease assessment of pediatric solid tumors remains a significant challenge, hindering clinical decision making and personalized treatment. Imaging-based techniques have inherent limitations, and children often require sedation, limiting their routine utility. In NBL, even disease specific tracer-based imaging like MIBG brings its own challenges, as not all tumors exhibit MIBG uptake, and differentiated tumors without neuroblastic cells can still exhibit mature neuronal components with tracer uptake and positive imaging findings. Non-invasive molecular monitoring with liquid biopsy can help solve these problems, yet commercially available sequencing based platforms fail to address this gap, as a significant proportion of HR-NBLs and pediatric solid tumors lack driver SNVs trackable in panel sequencing assays. There is a need for disease specific universal molecular markers that can be leveraged for global, accessible, and rapidly translatable liquid biopsy approaches.

In this study, we aimed to identify and leverage a robust NBL specific methylation-based biomarker panel to develop a mutation-agnostic metric that can support broad implementation of liquid biopsy-based disease assessment. We utilized ONT sequencing, which offers key advantages over conventional methylation platforms like the EPIC 850K array, reduced representation bisulfite sequencing (RRBS), or targeted capture panels. Unlike bisulfite-based methods requiring DNA conversion, ONT profiles native DNA methylation genome-wide while simultaneously capturing structural and sequence variants, critical for pediatric biopsies, where tissue is often limited. In addition, our whole-genome methylation sequencing approach captures clusters of CpG sites, rather than isolated CpGs, providing biological context and enhancing the interpretability of our findings. ONT long-read capability also facilitates analysis of extended cfDNA fragments, an emerging biomarker in liquid biopsies (24).

Global methylation profiling distinguished HR-NBL tumors from normal tissues, with PCA analysis revealing three molecular subgroups consistent with prior reports (19). *MYCN*-amplified tumors clustered with an 11q-deleted case in the “*MYCN*-type” group, while *ATRX* and *SMARCA4* mutated tumors (both SWI/SNF complex components) formed a distinct chromatin-remodeling subgroup, and an *ALK*-mutated tumor segregated separately. The co-segregation of tumors with *MYCN* amplification and 11q deletion has been shown previously (19), and may stem here from inclusion of a patient with rare co-occurrence of both *MYCN* amplification and 11q-deletion. While these genetic drivers classically represent exclusive molecular subtypes, co-segregation of their genome-wide methylation patterns may offer insights into convergent molecular pathways. These findings require validation in larger cohorts, and suggest the potential for complete molecular subtyping from a single diagnostic ONT run.

Integration of ONT-derived methylome maps with the normal human methylation atlas enabled the identification of 72 DMRs (meNBLs) relative to the atlas content. This atlas provides a high-resolution baseline of normal methylation signatures and may be considered as a “complex” control reference, representing a comprehensive collection of cell type-specific biomarkers from healthy origins, and enabling more accurate discrimination of NBL-specific methylation signals. As the atlas lacks fetal neural crest and its derivatives, and NBL arises from neural crest–derived sympathoadrenal precursor cells, we performed the analysis both with and without neuronal cell types to ultimately encompass both neuronal-origin and broader pan-cancer markers. By integrating the selected 25 meNBLs into the existing human methylome atlas (15), we extended the reference to include NBL as a distinct cell type, enabling the development of a mutation-agnostic metric for estimating NBL-derived DNA in cfDNA through deconvolution. The concordance of these markers in the TARGET EPIC array data and an independent validation cohort reflects the robustness of our biomarker selection, and findings will need to be validated in larger cohorts.

This approach successfully identified heterogeneous ranges of NBL-derived cfDNA in patient samples, while no signal was detected in cfDNA from healthy donors or from HR-NBL patients in remission. The simultaneous assessment of 25 meNBLs was found to be more accurate than IchorCNA estimation of disease burden based on available somatic mutation data (ddPCR and ONT sequencing), and it exhibits no background noise, enabling the reliable detection of NBL-derived cfDNA even at very low levels. In this context, other approaches such as the RASSF1A cfDNA methylation assay and cfRNA-based detection of TH and PHOX2B rely on single or dual markers, which may limit their sensitivity and specificity (25). Moreover, RASSF1A has been reported as a pan-tumor marker, rather than being specific to NBL (11). cfRNA-based assays also require specialized blood collection tubes that are not widely used in clinical practice, making them less practical to implement in routine diagnostic workflows compared to cfDNA-based methods. Furthermore, the methodology presented here for detecting meNBLs is readily adaptable for the identification of cancer-specific epigenetic markers in other malignancies. Beyond diagnosis, meNBLs may carry important mechanistic information not yet elucidated by genetic approaches, identifying novel DMRs with key roles in neurodevelopment, neuronal signaling and tumor–microenvironment interactions. Future work will explore the potential of the NBL methylome for drug target discovery.

In summary, we provide proof of concept that NBL-derived cfDNA methylation may provide a universal, mutation-agnostic metric for measuring disease burden across HR-NBL patients. Methylation-based cfDNA deconvolution provides accurate quantification of NBL cfDNA in the plasma, demonstrating the potential clinical utility for molecularly-defined relapse detection. These findings should be validated in larger cohorts, and highlight the potential role of cfDNA methylation-based liquid biopsy testing in directing patient care.

## Material and Methods

### Patient cohort and samples

Pediatric patient samples were obtained from individuals treated at the Schneider Children’s Medical Center of Israel between 2016-2024. The study was conducted in accordance with the Declaration of Helsinki and adhered to local guidelines and regulations. Ethical approval was obtained from the Institutional Review Board (IRB) of Rabin Medical Center (responsible for research conducted at Schneider Medical Center), under approval IRB# 4370-RMC. Written informed consent was obtained from participants or their legal representatives under approval number 0012-08-RMC. ONT sequencing was performed on seven HR-NBL tumor samples, twelve cfDNA samples from HR-NBL patients at diagnosis (five matched to the sequenced tumors and seven additional validation samples), along with cfDNA from eight healthy donors (Five adult and three pediatric samples) (Supplementary Table S1 and S2). Peripheral blood lymphocyte (PBL) pools, each comprising genomic DNA from ten individuals, served as additional control samples.

### DNA Isolation

Genomic DNA (gDNA) was extracted from fresh-frozen (FF) tumor tissue using spin columns (QIAamp DNA Micro Kit, Qiagen) according to the manufacturer’s instructions. The quantification of the isolated DNA was conducted using a Qubit Flex fluorometer with a 1x dsDNA HS Assay Kit (Invitrogen, Thermo Fisher). Whole blood was collected in EDTA Vacutainer tubes (BD Biosciences) and separated on the same day. Separation of plasma was performed by centrifugation for 10 minutes at 1600 g at room temperature, with the brake set to “off”, followed by a subsequent centrifugation step of 10 minutes at 1600 g at room temperature. The plasma was stored at −80°C until it was processed for cfDNA extraction. Isolation of cfDNA was performed starting from 400 µL to 3.5 mL of plasma. Smaller sample volumes were adjusted to 1 mL by adding 1x PBS (Sartorius).

After plasma was thawed and centrifuged for 10 minutes at 20,000 g at 4°C, cfDNA was extracted using the QIAamp Circulating Nucleic Acid Kit (Qiagen) with the Qiavac24s system, according to manufacturers’ recommendations. cfDNA concentration was measured using the Qubit Flex fluorometer with a 1x dsDNA HS Assay Kit (Invitrogen, Thermo Fisher). If cfDNA concentration was low but sufficient volume was available, samples were concentrated up to 2-fold using SpeedVac vacuum centrifugation. Size distribution of the cfDNA fragments was measured on a 4200 TapeStation system (Agilent Technologies, Santa Clara, CA).

### Library preparation and Nanopore Sequencing

Sequencing libraries were prepared using Ligation Sequencing Kit V14 (SQK-LSK114, Oxford Nanopore Technologies, UK) according to the manufacturers’ recommendations. cfDNA libraries were prepared using either Ligation Sequencing Kit V14 (SQK-LSK114) or Native Barcoding kit 24 V14 (SQK-NBD114.24), both from Oxford Nanopore Technologies, UK, according to the manufacturers’ recommendations. Sequencing was performed on a P2Solo device using R10.4.1 Flow cells (FLO-MIN114, Oxford Nanopore Technologies).

### Base-calling and Alignment

Base-calling and alignment were performed using ONT MinKNOW (version 24.06.16, Oxford Nanopore Technologies, UK), with the “modified base-calling” option enabled. The human reference genome hg38 was used for alignment. Reads from individual aligned BAM files were merged, sorted, and indexed using samtools (version 1.10). Since the reference atlas is based on bisulfite sequencing (which cannot distinguish 5mC from 5hmC), our Nanopore-detected 5hmC signals were converted to 5mC-equivalent using Modkit’s adjust-mods function prior to this analysis.

### DNA Methylation Analysis

DNA methylation beta values, representing methylation levels (0 = unmethylated to 1 = fully methylated), were calculated per CpG site using Modkit (version: 0.1.12, Oxford Nanopore Technologies, UK), with the --combine-strands and --cpg flags. Data analysis, including Principal Component Analysis (PCA), hierarchical clustering, correlation heatmaps, and violin plots, was performed in Python 3.10.7 using the pandas, scikit-learn, matplotlib, and seaborn libraries. All visualizations were generated with custom scripts (provided in the Supplementary Materials).

### NBL-Specific Marker Identification and Selection

Marker identification was based on the framework and reference data described by Loyfer et al. (15). Briefly, BAM files were converted to “beta” and “pat” formats using wgbstools with the bam2pat option and the --nanopore flag (15). The resulting PAT files (one per tumor DNA sample) were used in conjunction with a tissue-specific methylation dataset (GSE186458), which comprises 206 samples from 36 available different tissues. To identify uniformly methylated blocks, we applied the wgbstools segment option with parameters --min_cpg 3 --max_bp 10000, resulting in ∼4.3M blocks with a mean of 180bp and median of 379bp. This script detects genomic segments that exhibit consistent CpG methylation levels across all samples, including both our NBL tumors and the external reference dataset.

Differential methylation analysis was performed using the wgbstools find_markers command with default parameters. NBL samples were compared against all other tissues in the methylation dataset. In Loyfer et al. study, 3–4 replicates per cell type were sufficient to infer robust methylation patterns for biomarker identification. The inclusion of seven NBL tumors in our study aligns with their methodology and provides adequate representation. Given that NBL arises from neural crest cells, we repeated the analysis excluding neuronal cell types from the reference atlas, identifying additional DMRs associated with neuronal origin. Most regions were shared between the analyses including and excluding neuronal tissues (Supplementary Figure S1). Regions overlapping centromeres were excluded using bedtools intersect with GRCh38 centromere annotations (GRCh38.GCA_000001405.2_centromere_acen).

From an initial set of 188 DMRs, we first selected 76 regions exhibiting low mean methylation in healthy cfDNA (<10%). We then excluded two DMRs with high methylation in PBL pools (>20%), as gDNA contamination in cfDNA can elevate overall methylation levels. Two additional DMRs were manually excluded: one lacking sufficient coverage in ≥5/12 NBL cfDNA samples, and the other featuring a small CpG cluster (2 CpGs within a locus <80 bp). This analysis yielded a final set of 72 DMRs termed meNBLs. Methylation score was calculated using bedtools “map” with the -c mean flag to obtain an average value for all the different CpGs in the marker region. This was also done for public data obtained from the TARGET project (50).

The Genomic Regions Enrichment of Annotations Tool (GREAT) was used to affiliate markers to genes based on regulatory domains and genomic proximity. Gene Ontology (GO) enrichment analysis was carried out on the filtered subset of our DMRs. (Supplementary Figure S2). To enable comparison with methylation profiles from an independent ONT sequencing dataset, only tumors from patients younger than 30 years were included to better approximate the age distribution of our pediatric cohort. We further supplemented with adult breast and lung cancer samples (Supplementary Table S9).

### Deconvolution

25 selected meNBLs were added to the existing atlas (15) using uxmtools “build” option with the --use_um and --rlen 4 options to generate a modified atlas including the NBL entity. Cell of origin deconvolution was performed using uxmtools deconv option with our novel NBL atlas.

To ensure balanced representation of the different tissues in the modified atlas, which originally contained 25 markers per-tissue type, we selected a subset of 25 markers from the 72 meNBL markers. The selection maintained proportions between neuronal and non-neuronal features, mirroring the initial identification of 188 DMRs. For non-neuronal markers, selection was based on a multi-criteria ranking approach. Candidate DMRs were evaluated across five criteria: (1) highest CpG cluster span indicative of biologically meaningful regulatory regions, (2) lowest mean methylation in healthy control cfDNA, (3) lowest mean methylation in other advanced cancers, (4) highest mean methylation difference between tumor and healthy cfDNA, and (5) promoter-proximal location (±5 kb). Top-ranking candidates from the upper quartile of each criterion, especially those overlapping with multiple criteria, were selected for inclusion. Neuronal markers were prioritized based on brain tissue differential expression (GTEx database) and promoter location, ensuring a heterogeneous and balanced panel.

### Genetic Analysis

Single nucleotide variants (SNVs) and structural variants (SVs) were identified from whole-genome sequencing data across 60 genes selected through a comprehensive review of published genomic datasets on high-risk neuroblastoma (HR-NBL), as detailed in Supplementary Table S10. Variant calling was performed using Clair3 and Sniffles2 via the EPI2ME Labs workflow wf-human-variation. SNPs were subsequently annotated using ClinVar. Given the limited availability of comprehensive SV annotation databases for ONT data, a healthy control SV reference was constructed using peripheral blood lymphocyte (PBL) samples. SVs were called in the PBL samples using the same pipeline, merged using bcftools (version 1.3.1), and treated as a non-pathogenic reference set. Only SVs unique to the tumor samples (i.e., not overlapping with the control SVs) were retained for downstream interpretation. Copy number variations (CNVs) were detected using ichorCNA, applying 500 kbp bin sizes and default settings (23).

### Droplet Digital PCR

The QX200 ddPCR System (Bio-Rad Laboratories, CA, USA) was used to determine genomic aberrations in NB tissue/cell line gDNA or cfDNA. The ALK R1275Q (exon 25, position 3824, G>A) mutation was detected using Bio-Rad wet-lab assays dHsaCP250606768 and dHsaCP250606769. Analysis of the copy number statuses of MYCN (2p24.3) or ALK (2p23.2 to 2p23.1) was performed with two reference genes, NAGK (2p13.3), and AFF3 (2q11.2) in triplex ddPCR assays as described (10). PCR reactions were performed on a C1000 thermocycler (Bio-Rad Laboratories, CA, USA), with the annealing temperature of 55°C for ALK R1275Q mutation detection assay and 58°C for MYCN/ALK copy number assays as described (10). All ddPCR assays contained appropriate non-template, positive and negative controls in each run. Droplet reactions were assessed in the QX200 ddPCR Droplet Reader. Assays were analyzed using QX Manager Software version 2.2 (Bio-Rad Laboratories, CA, USA).

### Cell Culture

Genomic DNA from NBL cell lines served as positive and negative controls in ddPCR assays. The SK-N-AS and SH-SY5Y cell lines were purchased from ATCC (American Type Culture Collection, Manassas, VA, USA). LAN5 cell line were kindly gifted by Issac P. Witz and Orit Sagi-Assif. All cell lines were maintained at 37°C and 5% CO2, and grown in short-term culture. SK-N-AS cell lines were cultured in Dulbecco’s modified Eagle’s medium (Gibco, Life Technologies, UK) supplemented with 10% fetal calf serum and 1% nonessential amino acids. SH-SY5Y cell line were cultured in Dulbecco’s modified Eagle’s medium (Gibco, Life Technologies, UK) supplemented with 15-20% fetal calf serum. The LAN5 cell line was cultured in RPMI 1640 medium (Gibco, Life Technologies, UK) supplemented with 20% fetal calf serum. Cell lines were authenticated by high-throughput single-nucleotide polymorphism based assays (Hylabs, Rehovot Israel). For mutation detection ddPCR assays gDNA from cell lines was fragmented by sonication (Covaris M220).

### NBL panel sequencing

Tumor profiling was performed using the pan-pediatric cancer panel, Oncomine™ Childhood Cancer Research Assay (Ion Torrent, ThermoFisher) according to the manufacturer’s protocol. This tool analyzes the mutational status of 200 genes, including 82 mutation hotspots, 24 CNV targets, 44 genes with full exome coverage, and an RNA fusion panel of 97 genes. DNA and RNA libraries were generated using Ion AmpliSeq Library Preparation on the Ion Chef System (Thermo Fisher). Complementary DNA (cDNA) synthesis prior to library preparation for the RNA panel was carried out using SuperScript™ VILO™ Reverse Transcriptase (Thermo Fisher). Sequencing was performed on the Ion Torrent S5 (Thermo Fisher). Variants were identified and annotated by the Thermo Fisher Ion Reporter. A variant was accepted for further analysis if its coverage was at least 500 reads and the variant allele frequency (VAF) was greater than 5% (or 2% for hot spots). Amplification was reported when CN was above 8 at the 5% confidence level. Fusion genes were recognized as such if they had at least 10,000 per million reads. The identified variants were evaluated by different databases to analyze the clinical significance.

### Single-nucleotide polymorphism (SNP) array

Array analysis was performed using Affymetrix CytoScan HD array (Affymetrix, CA, USA) according to the manufacturer’s recommendations (Affymetrix manual protocol Affymetrix® Cytogenetics Copy Number Assay P/N 703038 Rev. 3). The raw data were processed using Chromosome Analysis Suite (ChAS) 4.5.0.34.

