## Supplementary_Figures for "Epigenetic Liquid Biopsy Enables Universal Mutation-Agnostic Molecular Surveillance for High-Risk Neuroblastoma"

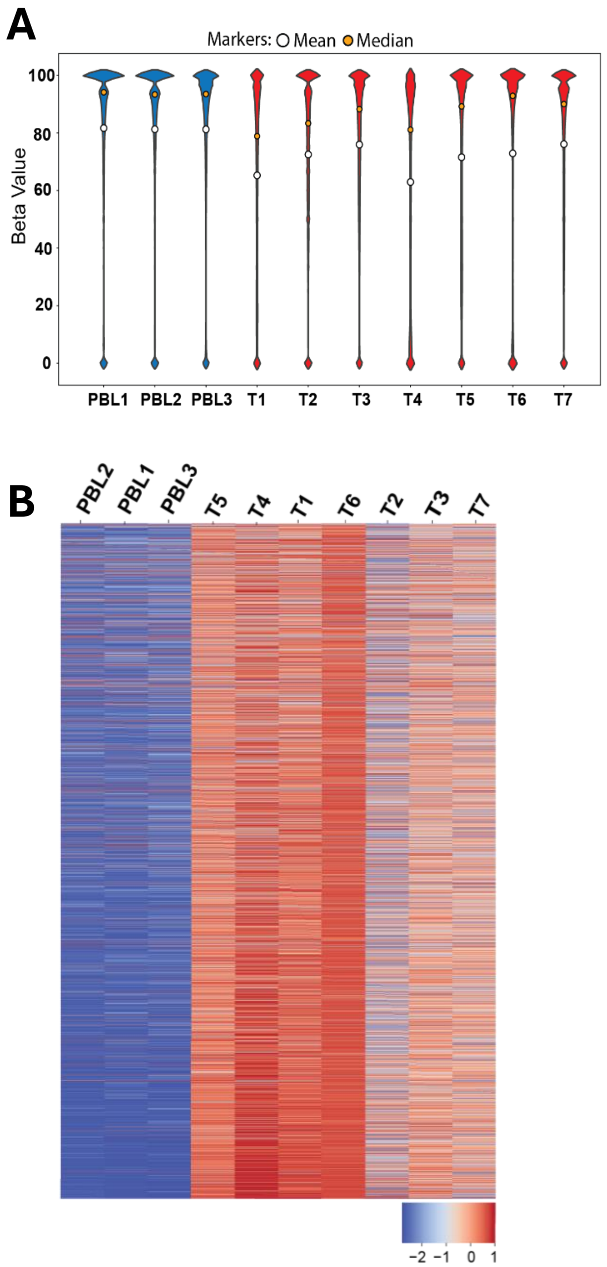


**Supplementary Figure S1**: DNA methylation analysis of NBL tumor samples reveals distinct methylation patterns. (A) Genome-wide DNA methylation levels were calculated in NBL 7 tumor gDNA and 3 PBL pools of 10 individuals each revealed globally reduced levels in NBL. Analysis is represented by violin plots with orange and white dots drawn at the distribution medians and means respectively. (B) Hierarchical clustering analysis based on methylation levels in NBL tumor and PBL pool samples reveal distinct molecular characteristics.

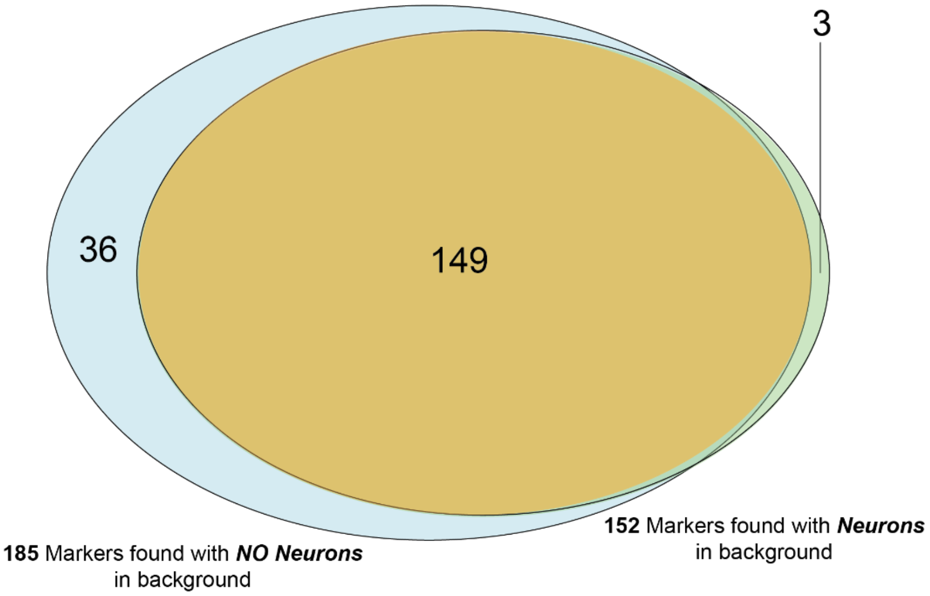


**Supplementary Figure S2**: Identification of NBL-specific DMRs through comparative methylome analysis to normal human cell-type atlas. Total of 152 NBL DMRs were identified in comparison to all tissue types in the human reference atlas (15). Given that NBL arises from neural crest cells, the analysis was repeated with neuronal cell types excluded from the atlas resulting in the identification of additional 36 "neuronal" regions.

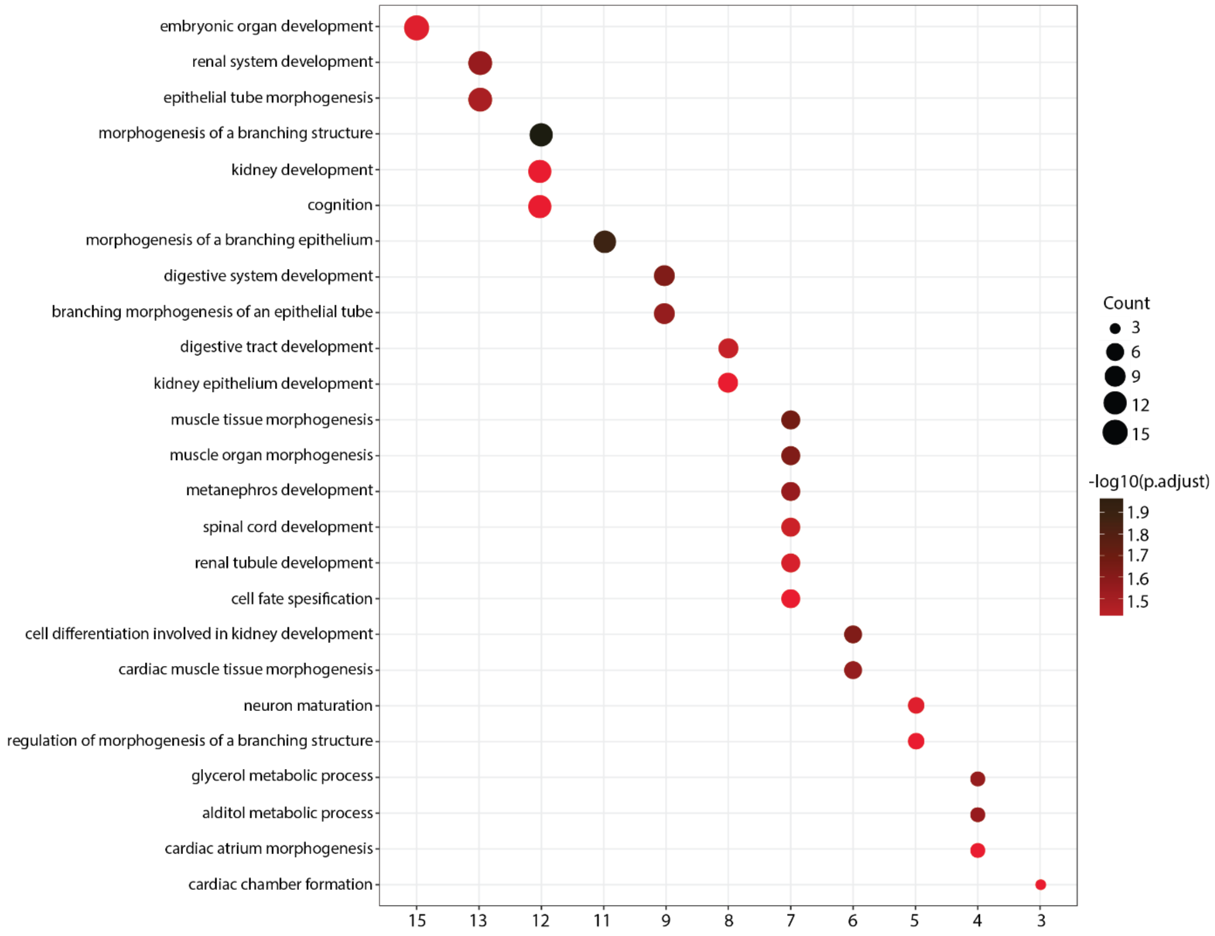


**Supplementary Figure S3**: Gene Ontology (GO) enrichment analysis of 152 identified DMRs. The x-axis represents the gene ratio (the proportion of genes from the methylated gene list involved in each GO term), while the size of the dots corresponds to the number of genes associated with each process. The color gradient indicates the adjusted p-value (−log10 p.adjust), reflecting the statistical significance of enrichment.

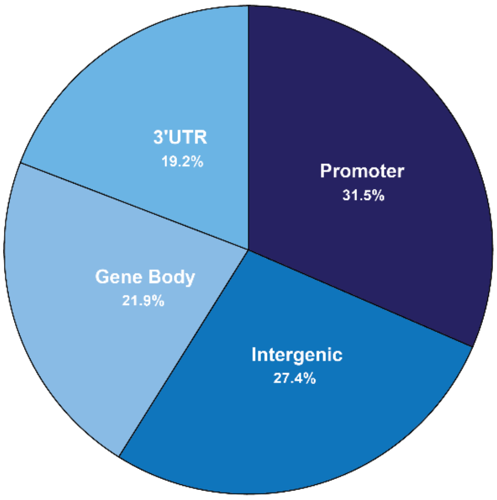


**Supplementary Figure S4:** Target distribution of 72 meNBLs across various genomic regions.

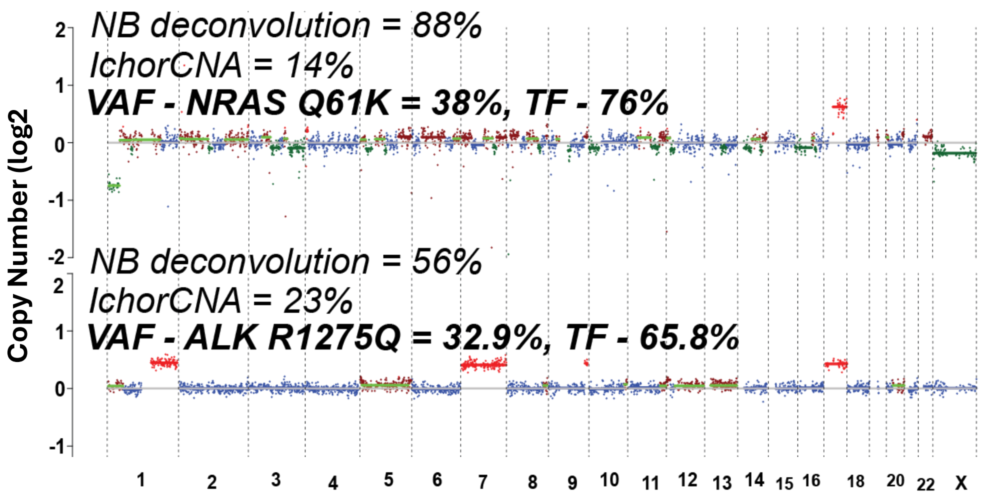


**Supplementary Figure S5**: Comparison of circulating NBL-deconvolution with IchorCNA tumor fraction estimates in two NBL cfDNA samples showing discrepancies
