## Supplementary material for "Epigenetic Liquid Biopsy Enables Universal Mutation-Agnostic Molecular Surveillance for High-Risk Neuroblastoma": Supplemetary_Tables1

### Supplementary Tables:

**Supplementary Table S1.** Clinical and Genomic Characteristics of NBL Tumor Samples Sequenced by ONT

| Sample | Age (years) | Sex | Disease course | MYCN amplification | TERT rearrangement | Genetic aberration | Typical CNVs | Coverage by ONT sequencing |
|---|---|---|---|---|---|---|---|---|
| T1_106897 | 5.7 | F | Death | No | | Not identified | 1p-, 3p-, 11q-, 17q+, +7 | 31.9 |
| T2_114998 | 2.4 | M | Death | No | | SMARCA4p.Gln194fs | 4p-, 11q-, 17q+ | 18.4 |
| T3_114596 | 5.2 | F | Death | No | | ALK p.Arg1275Gln | 2p+,5p+, 17q+, +7 | 38.7 |
| T4_217902 | 2.2 | F | Death | Yes | Yes | NRAS p.Gln61Lys | 1p-, 17q+ | 36.7 |
| T5_221663 | 11.6 | M | Death | Yes | | Not identified | 1p-, 11q-, 17q+, +7 | 34.6 |
| T6_204392 | 12.2 | M | Death | Yes | | TP53 p.Cys176Phe | 1p- | 32.1 |
| T7_202473 | 8.4 | M | No | No | Yes | ATRX p.Lys955Ter | 1p-,2p+, 4p-, 11q-, 17q+ | 28.4 |

**Supplementary Table S2:** Characteristics of cfDNA samples from NBL patients at diagnosis and controls analyzed by ONT sequencing

| | Sample | Molecular classification | Age (years) | Sex | Disease course | CNVs (Typical) | Oncomine Childhood Panel (Tumor) | ONT sequencing Coverage |
|---|---|---|---|---|---|---|---|---|
| Matched to Tumor NBL cfDNA | cfNB1_106697 | non MYCN | 5.7 | F | Death | 1p-, 3p-, 11q-, 17q+, +7 | Not identified | 3.08 |
| | cfNB2_115048 | non MYCN | 2.4 | M | Death | 4p-, 11q-, 17q+ | NA | 4.90 |
| | cfNB3_113096 | ALK p.Arg1275Gln | 5.2 | F | Death | 2p+,5p+, 17q+, +7 | ALK p.Arg1275Gln | 1.47 |
| | cfNB4_218072 | MYCN+TERT rearrangement | 2.2 | F | Death | 1p-, 17q+ | NRAS p.Gln61Lys | 12.98 |
| | cfNB5_215323 | MYCN | 11.6 | M | Death | 1p-, 11q-, 17q+, +7 | Not identified | 6.12 |
| Independent set NBL cfDNA | cfNB6_211373 | non MYCN | 1.9 | M | No | 1p-, 2p+, 3p-, 11q-, 17q+ | Not identified | 21.3 |
| | cfNB7_210823 | non MYCN | 2.8 | F | No | 1p-, 2p+, 4p-, 11q-, 17q+ | Not identified | 1.70 |
| | cfNB9_218011 | ALK p.Arg1275Gln | 10.1 | M | No | 1q+ | ALK p.Arg1275Gln | 1.05 |
| | cfNB10_214232 | MYCN | 6.6 | F | No | 1p-, 1q+ | NA | 0.64 |
| | cfNB11_216963 | MYCN+ALK amp | 2.4 | F | No | 1p-, 17q+ | Not identified | 7.52 |
| | cfNB12_202104 | MYCN+ALK p.Arg1275Gln | 3.4 | F | No | (-) | ALK p.Arg1275Gln | 2.27 |
| | cfNB13_107298 | MYCN | 8.7 | M | Death | 1p-, 17q+ | NA | 2.74 |
| Adult ctrl cfDNA | cfCtrl_Ad1 | NA | 41 | M | No | NA | NA | 2.9 |
| | cfCtrl_Ad2 | NA | 23 | F | No | NA | NA | 1.49 |
| | cfCtrl_Ad3 | NA | 25 | M | No | NA | NA | 3.4 |
| | cfCtrl_Ad4 | NA | 29 | M | No | NA | NA | 3.11 |
| | cfCtrl_Ad6 | NA | 32 | M | No | NA | NA | 8.76 |
| Pediatric Ctrl cfDNA | cfCtrlPd2 | NA | 13 | F | No | NA | NA | 0.45 |
| | cfCtrlPd3 | NA | 9 | M | No | NA | NA | 1.30 |
| | cfCtrlPd5 | NA | 11 | M | No | NA | NA | 1.67 |

**Supplementary Table S5:** List of 72 DMRs selected following comparative analysis with a normal human cell-type methylation atlas and cfDNA evaluation

| chr | start | end | Gene | Identification compared to reference atlas |
|---|---|---|---|---|
| chr1 | 34500637 | 34500709 | GJB5/C1or | Atlas wo Neurons |
| chr1 | 54800832 | 54801385 | LEXM /TTC | All atlas |
| chr1 | 2.05E+08 | 2.05E+08 | CDK18/LEI | All atlas |
| chr1 | 2.1E+08 | 2.1E+08 | TRAF3IP3/ | All atlas |
| chr1 | 2.1E+08 | 2.1E+08 | IRF6 | All atlas |
| chr1 | 2.1E+08 | 2.1E+08 | IRF6 | Atlas wo Neurons |
| chr2 | 44952303 | 44952738 | SIX3/SIX2 | All atlas |
| chr2 | 44952148 | 44952297 | SIX3/SIX2 | All atlas |
| chr2 | 44952754 | 44953051 | SIX3/SIX2 | All atlas |
| chr2 | 2.18E+08 | 2.18E+08 | PNKD/CAT | Atlas wo Neurons |
| chr2 | 2.19E+08 | 2.19E+08 | DES | All atlas |
| chr2 | 2.19E+08 | 2.19E+08 | DES | All atlas |
| chr3 | 1.4E+08 | 1.4E+08 | NMNAT3 | All atlas |
| chr3 | 1.4E+08 | 1.4E+08 | NMNAT3 | All atlas |
| chr3 | 1.58E+08 | 1.58E+08 | SHOX2/VE | All atlas |
| chr4 | 8858633 | 8859201 | HMX1/CPZ | All atlas |
| chr4 | 8859311 | 8859822 | HMX1/CPZ | All atlas |
| chr4 | 8857116 | 8857582 | HMX1/CPZ | All atlas |
| chr4 | 8857601 | 8858396 | HMX1/CPZ | All atlas |
| chr4 | 8877216 | 8878163 | HMX1/FAM | All atlas |
| chr5 | 72556517 | 72557083 | ZNF366/TN | All atlas |
| chr5 | 1.54E+08 | 1.54E+08 | HAND1 (-1 | All atlas |
| chr5 | 1.71E+08 | 1.71E+08 | TLX3/NPM | Atlas wo Neurons |
| chr6 | 26087378 | 26088004 | HFE | Atlas wo Neurons |
| chr6 | 1.12E+08 | 1.12E+08 | TRAF3IP2 | Atlas wo Neurons |
| chr6 | 1.57E+08 | 1.57E+08 | ARID1B/TN | All atlas |
| chr7 | 35257129 | 35257671 | TBX20 | All atlas |
| chr7 | 35261977 | 35262053 | TBX20 | All atlas |
| chr7 | 51591365 | 51591488 | COBL | Atlas wo Neurons |
| chr7 | 76282099 | 76283042 | SRRM3/HS | Atlas wo Neurons |
| chr7 | 1.27E+08 | 1.27E+08 | GRM8 | Atlas wo Neurons |
| chr7 | 1.28E+08 | 1.28E+08 | intergenic | All atlas |
| chr7 | 1.28E+08 | 1.28E+08 | PRRT4/RBI | All atlas |
| chr7 | 1.28E+08 | 1.28E+08 | PRRT4/RBI | Atlas wo Neurons |
| chr7 | 1.49E+08 | 1.49E+08 | ZNF777/ZN | Atlas wo Neurons |
| chr8 | 1701369 | 1701627 | DLGAP2 | Atlas wo Neurons |
| chr8 | 1701164 | 1701361 | DLGAP2 | Atlas wo Neurons |
| chr8 | 13131921 | 13132451 | DLC1/KIAA | All atlas |
| chr9 | 1.26E+08 | 1.26E+08 | MVB12B/P | All atlas |
| chr9 | 1.29E+08 | 1.29E+08 | NTMT1/PTI | All atlas |
| chr9 | 1.3E+08 | 1.3E+08 | C9ORF50/ | Atlas wo Neurons |
| chr9 | 1.38E+08 | 1.38E+08 | CACNA1B | All atlas |
| chr10 | 42932794 | 42933504 | RET/BMS1 | Atlas wo Neurons |
| chr11 | 2791633 | 2791921 | CDKN1C/K | All atlas |
| chr11 | 62923660 | 62923914 | CHRM1 | All atlas |
| chr12 | 1796681 | 1796870 | CACNA2D | All atlas |
| chr12 | 1.27E+08 | 1.27E+08 | intergenic | All atlas |
| chr12 | 49297294 | 49297921 | PRPH/TRO | All atlas |
| chr12 | 49296952 | 49297292 | PRPH/TRO | All atlas |
| chr13 | 42991938 | 42992483 | EPSTI1/DN | Atlas wo Neurons |
| chr14 | 89027078 | 89027628 | TTC8/FOX | All atlas |
| chr15 | 34795223 | 34795805 | ACTC1 | All atlas |
| chr15 | 89371299 | 89371908 | POLG/RHO | All atlas |
| chr15 | 89371150 | 89371286 | POLG/RHO | All atlas |
| chr15 | 89378604 | 89378854 | POLG/RHO | Atlas wo Neurons |
| chr15 | 89409285 | 89409402 | POLG/RHO | Atlas wo Neurons |
| chr16 | 695547 | 695893 | FBXL16/W | Atlas wo Neurons |
| chr16 | 89574777 | 89574962 | CPNE7 | Atlas wo Neurons |
| chr17 | 34638584 | 34638762 | TMEM132I | Atlas wo Neurons |
| chr17 | 36657720 | 36658246 | LHX1/MRM | Atlas wo Neurons |
| chr17 | 41527495 | 41528084 | KRT19 | All atlas |
| chr17 | 43006678 | 43006969 | IFI35 | Atlas wo Neurons |
| chr17 | 43952860 | 43953252 | PPY/PYY | All atlas |
| chr17 | 74212399 | 74212818 | TTYH2 | Atlas wo Neurons |
| chr19 | 1470223 | 1470547 | APC2/C19 | Atlas wo Neurons |
| chr19 | 1470555 | 1470908 | APC2/C19 | Atlas wo Neurons |
| chr19 | 1470923 | 1471242 | APC2/C19 | Atlas wo Neurons |
| chr19 | 13872735 | 13873374 | NANOS3 | Atlas wo Neurons |
| chr19 | 46413249 | 46413733 | CCDC8 | Atlas wo Neurons |
| chr20 | 2558569 | 2558813 | TMC2/NOI | All atlas |
| chr20 | 2558817 | 2559219 | TMC2/NOI | All atlas |
| chr20 | 59006754 | 59007419 | CTSZ | Atlas wo Neurons |

**Supplementary Table S6:** List of NBL-Specific DMRs for cfDNA deconvolution analysis

| Chr | Start | End | Name | Class |
|---|---|---|---|---|
| chr1 | 54800833 | 54801385 | LEXM /TTC22 | "Pan Cancer" |
| chr1 | 205430663 | 205431108 | CDK18/LEMD1/BLACAT1 | "Pan Cancer" |
| chr1 | 209747770 | 209748026 | TRAF3IP3/HSD11B1 | "Pan Cancer" |
| chr1 | 209806011 | 209806291 | IRF6 | "Pan Cancer" |
| chr3 | 139677637 | 139678126 | NMNAT3 | "Pan Cancer" |
| chr4 | 8877217 | 8878163 | HMX1/CPZ | "Pan Cancer" |
| chr6 | 157235646 | 157236222 | ARID1B/TMEM242 | "Pan Cancer" |
| chr7 | 35257130 | 35257671 | TBX20 | "Pan Cancer" |
| chr9 | 129383416 | 129383947 | NTMT1/PTPA | "Pan Cancer" |
| chr9 | 138023371 | 138023711 | CACNA1B | "Pan Cancer" |
| chr11 | 62923661 | 62923914 | CHRM1 | "Pan Cancer" |
| chr12 | 1796682 | 1796870 | CACNA2D4 /LRTM2/ADIPOR4 | "Pan Cancer" |
| chr12 | 49296953 | 49297292 | intergenic | "Pan Cancer" |
| chr14 | 89027079 | 89027628 | TTC8/FOXN3 | "Pan Cancer" |
| chr15 | 34795224 | 34795805 | ACTC1 | "Pan Cancer" |
| chr15 | 89371300 | 89371908 | POLG/RHCG/MIR9-3HG | "Pan Cancer" |
| chr16 | 695548 | 695893 | FBXL16/WDR24 | "Neuronal" |
| chr16 | 89574778 | 89574962 | CPNE7 | "Neuronal" |
| chr17 | 41527496 | 41528084 | KRT19 | "Pan Cancer" |
| chr17 | 43006679 | 43006969 | IFI35 | "Neuronal" |
| chr17 | 43952861 | 43953252 | PPY/PYY | "Pan Cancer" |
| chr19 | 1470924 | 1471242 | APC2/C19orf25 | "Neuronal" |
| chr19 | 46413250 | 46413733 | CCDC8 | "Neuronal" |
| chr20 | 2558570 | 2558813 | TMC2/NOP56 | "Pan Cancer" |
| chr20 | 59006755 | 59007419 | CTSZ | "Neuronal" |

**Supplementary Table S7:** NBL cfDNA quantification in patients and controls: Deconvolution vs ichorCNA

| | Sample | Molecular classification | NBL deconvolution % | iCoreCNA TF % |
|---|---|---|---|---|
| Matched to Tumor NBL cfDNA | cfNB1_106697 | non MYCN | 100 | 85.6 |
| | cfNB2_115048 | non MYCN | 0.43 | 9.8 |
| | cfNB3_113096 | ALK mutation | 73 | 54.2 |
| | cfNB4_218072 | MYCN+TERT rearrangement | 87.9 | 14.0 |
| | cfNB5_215323 | MYCN | 100 | 97.4 |
| Independent set NBL cfDNA | cfNB6_211373 | non MYCN | 23.1 | 61.7 |
| | cfNB7_210823 | non MYCN | 100 | 55.7 |
| | cfNB9_218011 | ALK mutation | 55.7 | 22.8 |
| | cfNB10_214232 | MYCN | 66.7 | 13.9 |
| | cfNB11_216963 | MYCN+ALK amp | 73 | 59.9 |
| | cfNB12_202104 | MYCN+ALK mutation | 100 | 58.0 |
| | cfNB13_107298 | MYCN | 90 | 29.6 |
| Adult ctrl cfDNA | cfCtrl_Ad1 | NA | 0 | 6.35 |
| | cfCtrl_Ad2 | NA | 0 | 6.27 |
| | cfCtrl_Ad3 | NA | 0 | 9.9 |
| | cfCtrl_Ad4 | NA | 0 | 10.88 |
| | cfCtrl_Ad6 | NA | 0 | 8.52 |
| Pediatri c Ctrl cfDNA | cfCtrlPd2 | NA | 0 | NA |
| | cfCtrlPd3 | NA | 0 | 10.08 |
| | cfCtrlPd5 | NA | 0 | 11.25 |

**Supplementary Table S8:** List of 60 NBL-associated genes from published studies analyzed for genomic aberrations in seven NBL tumors

| Gene | Chr | Start | End |
|---|---|---|---|
| AKT3 | chr1 | 243651536 | 244014381 |
| ARID1A | chr1 | 27022507 | 27108595 |
| MDM4 | chr1 | 204485535 | 204527248 |
| NRAS | chr1 | 115247091 | 115259392 |
| ALK | chr2 | 29415641 | 30144452 |
| DNMT3A | chr2 | 25450744 | 25565459 |
| MYCN | chr2 | 16075672 | 16092126 |
| ACVR2A | chr2 | 148597087 | 148693391 |
| ATR | chr3 | 142168078 | 142297575 |
| BCL6 | chr3 | 187439166 | 187463256 |
| CTNNB1 | chr3 | 41240997 | 41281934 |
| PIK3CA | chr3 | 178866146 | 178957881 |
| SETD2 | chr3 | 47052926 | 47210603 |
| PDGFRA | chr4 | 55090460 | 55169412 |
| PDGFRA | chr4 | 55090460 | 55169412 |
| PHOX2B | chr4 | 41746100 | 41750742 |
| FGFR4 | chr5 | 176513917 | 176525145 |
| TERT | chr5 | 1253283 | 1295183 |
| ARID1B | chr6 | 157097161 | 157531913 |
| CDKN1A | chr6 | 36641491 | 36660109 |
| DAXX | chr6 | 33286335 | 33290736 |
| LIN28B | chr6 | 105399982 | 105536207 |
| ROS1 | chr6 | 117603516 | 117752105 |
| BRAF | chr7 | 140413129 | 140624729 |
| MET | chr7 | 116307250 | 116443431 |
| FGFR1 | chr8 | 38263661 | 38331153 |
| MYC | chr8 | 128743477 | 128760197 |
| STMN2 | chr8 | 80518352 | 80583393 |
| CDKN2A | chr9 | 21962751 | 21999391 |
| CDKN2B | chr9 | 22002903 | 22009312 |
| PTCH1 | chr9 | 98200262 | 98284253 |
| PTPRD | chr9 | 8314247 | 10613002 |
| TSC1 | chr9 | 135766737 | 135820003 |
| ARID5B | chr10 | 63661459 | 63856703 |
| PTEN | chr10 | 89618382 | 89736687 |
| ATM | chr11 | 108088794 | 108244829 |
| CCND1 | chr11 | 69455925 | 69469242 |
| HRAS | chr11 | 527242 | 540576 |
| CDK4 | chr12 | 58141511 | 58146093 |
| MDM2 | chr12 | 69201953 | 69244466 |
| PTPN11 | chr12 | 112851751 | 112952722 |
| BRCA2 | chr13 | 32889646 | 32974405 |
| RB1 | chr13 | 48877888 | 49056026 |
| AKT1 | chr14 | 105235687 | 105262085 |
| DICER1 | chr14 | 95547565 | 95628866 |
| BRD7 | chr16 | 50349869 | 50402899 |
| CREBBP | chr16 | 3770055 | 3935714 |
| TSC2 | chr16 | 2092986 | 2144492 |
| CDK12 | chr17 | 37617740 | 37690797 |
| ERBB2 | chr17 | 37851317 | 37889911 |
| NF1 | chr17 | 29421946 | 29704695 |
| SMARCE1 | chr17 | 38781215 | 38804070 |
| TP53 | chr17 | 7566739 | 7595808 |
| ALPK2 | chr18 | 56148480 | 56296323 |
| AKT2 | chr19 | 40736225 | 40791252 |
| JAK3 | chr19 | 17935592 | 17958791 |
| SMARCA4 | chr19 | 11066706 | 11177949 |
| MAPK1 | chr22 | 22108946 | 22226935 |
| ATRX | chrX | 76760359 | 77041702 |
| BCOR | chrX | 39910500 | 40036582 |

**Supplementary Table S10:** cfDNA Samples from NBL patients at relapse and remission for deconvolution-based disease quantification

| Sample | MYCN/non MYCN | Age (years) | SEX | Disease Course | NBL deconvolution % | IChoreCNA TF% | ONT Coverage |
|---|---|---|---|---|---|---|---|
| cfNB_107298 | MYCN | 8.7 | M | At diagnosis | 90.0 | 29.6 | 2.7 |
| cfNB_116308 | MYCN | 9.1 | M | no disease | 0.0 | 7.7 | 4.3 |
| cfNB_105139 | MYCN | 9.5 | M | no disease | 0.0 | 8.7 | 1.7 |
| cfNB_110849 | MYCN | 9.8 | M | Progression | 40.6 | 31.8 | 4.5 |
| cfNB_115029 | MYCN | 10.1 | M | Progression | 88.0 | 80.1 | 5.2 |
| cfNB_112968 | no MYCN | 6.9 | M | At diagnosis | 18.0 | 26.6 | 3.1 |
| cfNB_113319 | no MYCN | 7.9 | M | no disease | 0.0 | 10.5 | 7.4 |
| cfNB_118459 | no MYCN | 8.1 | M | no disease | 0.0 | 8.1 | 2.7 |
| cfNB_205010 | no MYCN | 9.5 | M | Progression | 5.0 | 12.8 | 1.1 |
| cfNB_109256 | MYCN | 2.3 | M | no disease | 0.0 | 9.3 | 1.3 |
| cfNB_113077 | MYCN | 3.3 | M | no disease | 0.0 | 7.5 | 1.0 |
| cfNB_106148 | MYCN | 4.0 | M | Progression | 87.0 | 39.6 | 3.8 |
| cfNB_218613 | MYCN | 13.9 | M | no disease | 0.0 | 12.2 | 0.8 |
| cfNB_200214 | MYCN | 14.0 | M | Progression | 100.0 | 47.0 | 1.9 |
| cfNB_209025 | nonMYC | 6.4 | M | Progression | 86.5 | 85.0 | 1.8 |
| cfNB_207434 | nonMYC | 6.2 | M | Progression | 86.1 | 42.9 | 0.37 |
